# Chromosome-level genome assemblies of five *Sinocyclocheilus* species

**DOI:** 10.1101/2025.02.27.640546

**Authors:** Chao Bian, Ruihan Li, Yuqian Ouyang, Junxing Yang, Xidong Mu, Qiong Shi

## Abstract

*Sinocyclocheilus*, a genus of tetraploid fishes, is endemic to the karst regions of Southwest China. All species within this genus are classified as second-class national protected species due to their unique and fragile habitat. However, absence of high-quality genomic resources has hindered various research efforts to elucidate their phylogenetic relationships and the origin of polyploidy. To address these academic challenges, we at first constructed a high-quality genome assembly for the most abundant representative, golden-line barbel (*Sinocyclocheilus grahami*), by integration of PacBio long-read and Hi-C sequencing technologies. The final scaffold-level genome assembly of *S. grahami* is 1.6 Gb in length, with a scaffold N50 up to 30.7 Mb. A total of 42,205 protein-coding genes were annotated. Subsequently, 93.1% of the assembled genome sequences (about 1.5 Gb) and 93.8% of the total predicted genes were successfully anchored onto 48 chromosomes. Furthermore, we obtained chromosome-level genome assemblies for four other *Sinocyclocheilus* species (including *S. anophthalmus, S. maitianheensis, S. anshuiensis*, and *S. rhinocerous*) based on homologous comparison. These genomic data we present in this study provide valuable genetic resources for in-depth investigation on cave adaptation and improvement of economic values and conservation of diverse *Sinocyclocheilus* fishes.

## INTRODUCTION

*Sinocyclocheilus* (order: Cypriniformes; family: Cyprinidae; subfamily: Barbinae) is a genus of tetraploid fishes endemic to the karst regions of the Yunnan-Guizhou plateau and surrounding areas in Southwest China, including Guangxi, Guizhou, Yunnan, and Hubei provinces [1]. All members from this genus are classified as second-class protected species, highlighting the urgent need for their conservation and further investigation. Despite recent R & D efforts, such as an artificial breeding program for *S. yunnanensis* to prevent extinction [2], many other species, particularly those with small amount and limited distributions, remain in a threatened status.

Due to the long-term geographic isolation, *Sinocyclocheilus* species have undergone significant speciation, making it the most species-rich genus of cavefish, with 76 known members [1]. This genus resides in various ecological environments, ranging from surface-dwelling, semi-cave-dwelling and cave-restricted. These distinct habitat types result in trait diversity in morphology, behavior, and physiology [3], making them a good model for studying cave adaptation and phylogenetic evolution. Although *Sinocyclocheilus* is of significant scientific interests, high-quality genomic resources and whole genome-based comparative studies are rare among diverse *Sinocyclocheilus* fishes. The lack of genomic information impedes deep understanding of key evolutionary questions, such as phylogenetic relationships, the origin of polyploidy, and the evolution of ancestral chromosomes within this genus.

To enrich the genetic resources for *Sinocyclocheilus* members, we at first constructed a chromosome-level genome assembly for the most abundant and surface-dwelling representative, *S. grahami* (locally named as golden-line barbel) using PacBio and Hi-C sequencing technologies, and subsequently conducted homologous comparison to obtain chromosome-level genome assemblies for other four *Sinocyclocheilus* species (including surface-dwelling *S. maitianheensis*, semi-cave-dwelling *S. rhinocerous*, and cave-restricted *S. anshuiensis & S. anophthalmus*). To verify the allotetraploid origin of *Sinocyclocheilus*, we conducted a genome-wide synteny analysis between *S. grahami* and its relative, common carp (*Cyprinus carpio*). The analysis revealed extensive chromosomal rearrangements and supported the allotetraploid origin of the *Sinocyclocheilus* genus. These genomic data we present in this paper provide valuable genetic resource for deeply investigating cave adaptation mechanisms and exploring potential economic and ecological values of diverse *Sinocyclocheilus* species.

## METHODS

### Sample collection, DNA extraction, and genome and transcriptome sequencing

Muscle sample of artificially bred *S. grahami* was collected from the Endangered Fish Conservation Center of Kunming Institute of Zoology, which is located in Kunming city, Yunnan province, China. Genomic DNA (gDNA) and total RNA were extracted by using a Nucleic Acid Kit (Qiagen, Germantown, MD, USA) and TRIZOL Reagent (Invitrogen, Carlsbad, CA, USA) respectively following the manufacturers’ instructions.

Multiple sequencing strategies were applied for construction of a whole-genome assembly of *S. grahami*. In brief, the draft genome assembly based on Illumina sequencing technology (Illumina Inc., San Diego, CA, USA) was obtained from our previous study [4] as a reference. The genomic DNA from muscle in our present study was used to construct a SMART bell library with an insert size of 20 kb, and this library was subsequently sequenced on a PacBio Sequel platform (Pacific Biosciences, Menlo Park, CA, USA). For construction of a chromosome-level genome assembly, a Hi-C (High-through chromosome conformation capture) library was generated for sequencing on an Illumina HiSeq X-Ten platform. In addition, a paired-end library with an insert size of 400 bp was constructed from the extracted gDNA and then sequenced on an Illumina HiSeq X-Ten platform for genome size estimation. Adapters, duplicated reads and low-quality reads with 10 or more N bases were removed by SOAPfilter v2.2 [5]. For transcriptome sequencing (to support annotation of genes), a paired-end library with an insert size of 350 bp was generated and then sequenced on an Illumina HiSeq X-Ten platform. Raw data were filtered by SOAPnuke v1.0 [6].

### Genome size estimation and chromosome-level genome assembly

A 17-mer frequency distribution, which was confirmed to be a Poisson pattern [7], was applied to estimate the genome size of *S. grahami* with the library of short-inserted size (400 bp). The genome size calculation formula is set as follows [4]: Genome Size=Knum/Kdepth (Knum is the number of 17-mer; Kdepth is the sequencing depth at the core peak frequency).

Based on our published contigs of *S. graham* [4], we performed a hybrid genome assembly by combination of short contigs [4] with PacBio long reads (this study) into the primary scaffolds by using DBG2OLC v1.1 [8] with defaulted parameters. These scaffolds were subsequently extended using SSPACE v2.0 [9]. Minimap [10] and Racon [11] were employed for two rounds of error correction to obtain the final scaffolds with assistance of PacBio long reads.

The Hi-C raw reads were mapping onto these scaffolds by Bowtie2 [12] and quality control by HiC-Pro v2.8.0 [13] to obtain valid data for genome-wide interaction matrix construction. Juicer v1.5 [14] and 3D-DNA *de novo* v170123 [15] were applied to arrange and orientate scaffolds into chromosomes. A Hi-C heatmap was drawn by HiCPlotter v0.6.6 [16] for visualization.

### Repeat annotation, gene prediction, and function prediction

Three prediction methods were combined for repeat annotation, including *de novo*, homolog-based and tandem repeat prediction. A *de novo* repeat library was built by RepeatModeler v1.04 [17] and LTR_FINDER v1.0.6 [18]. Genome sequences were mapped onto this library to identify repeat sequences using RepeatMasker v4.06 [19]. For the homolog-based prediction, transposable elements (TEs) were identified using RepeatMasker v4.06 and RepeatProteinMask v4.06 [19] based on the RepBase TE v21.01 [20] library. Tandem repeat sequences were final identified by Tandem Repeat Finder v4.09 [21].

Protein-coding genes were predicted by integration of three methods, including homology-based annotation, *de novo* prediction and transcriptome-based annotation. Protein sequences of five representative teleost species were downloaded from NCBI GenBank [22] for genome-wide mapping onto *S. grahami*, including zebrafish (*Danio rerio*, NCBI accession: GCF_000002035.6), medaka (*Oryzias latipes*, NCBI accession: GCF_002234675.1), *S. anshuiensis, S. rhinocerous* and *S. grahami* (the primary genome assemblies with Illumina data). BLAT [23] and GeneWise v2.4.2 [24] were applied for sequence alignment and gene structure prediction. Augustus v3.2.1

[25] tool was used to *de novo* predict coding sequences (CDS) after the repeat elements were masked. Hisat v0.1.6 [26] and Cufflink v2.2.1 [27] was employed to perform the transcriptome-based annotation. Finally, a non-redundant gene set was merged by MAKER v2.31.8 [28]. For function annotation, we searched four public databases (Swiss-Prot [29], TrEMBL [29], Interpro [30] and KEGG [31]) as references to complete the annotation of gene functions.

### Pseudochromosome construction of another four scaffold-level assemblies of different *Sinocyclocheilus* fishes

The general chromosome number of *Sinocyclocheilus* fishes is 96 [32]. Pairwise whole-genome alignments were employed to construct pseudochromosomes of scaffold-level assemblies for four other *Sinocyclocheilus* fishes (*S. anophthalmus, S. maitianheensis, S. anshuiensis*, and *S. rhinocerous*) based on the reference chromosome-level assembly of *S. grahami*. Lastz v1.1 [33] was used to process the genome alignments. Those aligned sequences with a length of more than 10 kb were retained for pseudochromosome construction. Synteny blocks of each genome for the total five *Sinocyclocheilus* members were identified using MCScanX [34] respectively, after self-aligning with its own protein set by BLAST [35] using the optimized parameter of E-value less than 1e-5. Circos figures were plotted by the Circos software [36].

### Subgenome identification in *S. grahami* and phylogenetic analysis

Common carp (*Cyprinus carpio*) is a well-known allotetraploid species [37] that shared the recent genome-wide duplication event with *Sinocyclocheilus* species as we reported earlier [38]. Subgenomes A and B of Common carp [39] were used as the references to identify corresponding synteny blocks in goldfish (*Carassius auratus*), *S. grahami* and *S. anophthalmus* genomes for subsequent subgenomes construction by using MUMmer v4.0beta1[40]. RectChr (https://github.com/BGI-shenzhen/RectChr) was applied to visualize the synteny blocks and chromosome structure variations.

For a phylogenetic analysis, BLASTp [35] and OrthoMCL [41] were performed for protein sequence alignment and gene family clustering. All the single-copy orthologous genes were aligned by using MUSCLE v3.8.31 [42] for all genomes and subgenomes. Subsequently, Gblocks [43] was used to obtain the conservative multi-sequence alignments. Finally, we employed PhyML v3.0 [44] to construct a phylogenetic tree using the maximum likelihood method. MCMCtree in the PAML package [45] was employed to estimate the divergence time from above-mentioned fishes and other representative species.

## RESULTS AND DISCUSSION

### Chromosome-level genome assemblies of the five *Sinocyclocheilus* Species

A total of 86.4 Gb, 79.1 Gb and 229.0 Gb of Illumina, Pacbio and Hi-C reads were sequenced. We constructed a chromosome-level genome assembly for *S. grahami* using PacBio and Hi-C sequencing technologies. For the K-mer analysis, we estimate the genome size is 1.9 Gb. The final chromosome-level genome assembly of *S. grahami* is 1.6 Gb with a contig N50 of 738.5 kb and a scaffold N50 of 30.7 Mb.

About 93.1% of the assembled genome sequences (1.5 Gb) and 93.8% of the predicted genes were anchored onto 48 chromosomes (**Figure 1A-B**).

**Figure 1.**
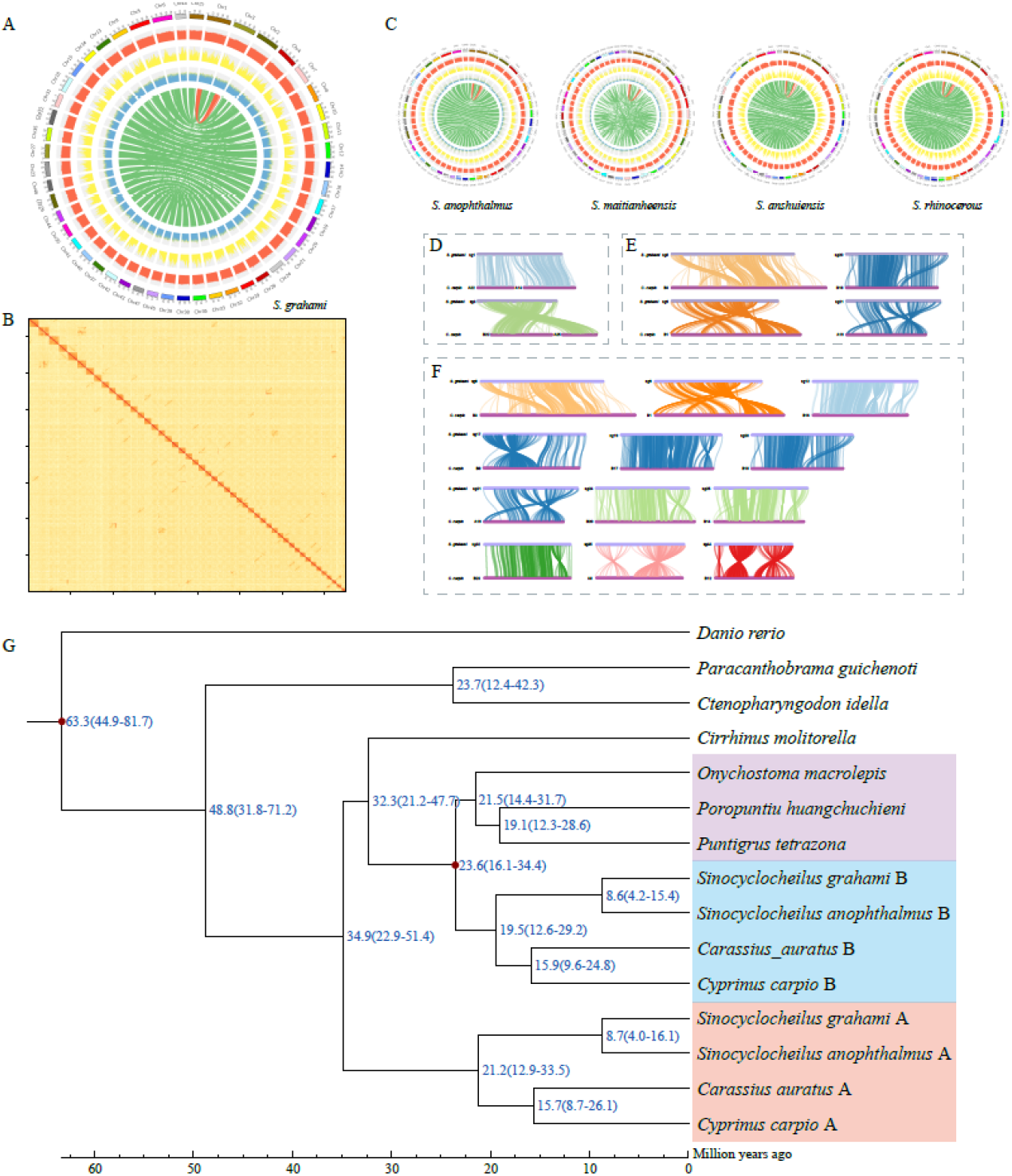
(A) Circos atlas of the reference chromosome-level genome assembly of *S. grahami*. Rings from outside to inside include chromosome length (Mb), distribution of gene density in each 100-kb non-overlapping genomic window, distribution of SNP density in each 100-kb non-overlapping genomic window, GC content in each 100-kb non-overlapping genomic window, and internal syntenic blocks of chromosomes that were connected by green lines. Red lines mark a special syntenic block between Chr1 and Chr3. (B) Genome-wide Hi-C heatmap of the *S. grahami* genome. (C) Circos atlases of the chromosome-level genome assemblies of four other *Sinocyclocheilus* species. (D-F) Two chromosomal fusionsn, five chromosomal translocations and eighteen chromosomal inversion events between *S. grahami* (top) and *C. carpio* (bottom). (G) A phylogenetic tree of seven vertebrate genomes and eight sub-genomes of tetraploid species. Orange box represents the clade of sub-genome A; blue box marks the clade of sub-genome B; purple box highlights a clade homologous to the ancestors of sub-genome A. Diverge time is numbered in blue, and a geographic time scale in million years ago (Mya) is provided.

A total of 583.2 Mb repeat sequences were annotated (**Table 1**). A sum of 42,205 protein-coding genes were annotated from the *S. grahami* genome assembly (**Table 2**), and 39,458 (93.5% of all) genes were annotated with functions. We also constructed chromosome-level genome assemblies for other four *Sinocyclocheilus* species based on homologous comparison (**Figure 1C**). Over 82% of the genome sequences of all the four species were anchored on these constructed chromosomes.

**Table 1.** Statistics of repeat sequences among the *S. grahami* genomes.

| Type | *S. grahami* |  |
| --- | --- | --- |
|  | Repeat Size (bp) | % of genome |
| ProteinMask | 72,493,669 | 4.6 |
| RepeatMasker | 330,527,709 | 20.8 |
| TRF | 33,729,084 | 2.1 |
| *De novo* | 372,094,752 | 23.4 |
| Total | 583,165,599 | 36.7 |
| DNA | 374,708,178 | 23.6 |
| LINE | 105,180,165 | 6.6 |
| SINE | 5,429,507 | 0.3 |
| LTR | 145,884,791 | 9.2 |
| Other | 4,064 | 0 |
| Unknown | 2,578,927 | 0.2 |
| Total | 547,757,670 | 34.5 |

**Table 2.** Protein-coding gene annotation of *S. grahami* genome.

| Method | Software or Speceis | Gene number | Average |  |  |  |  |
| --- | --- | --- | --- | --- | --- | --- | --- |
|  |  |  | Transcript Length ( bp ) | CDS Length ( bp ) | Exons per Gene | Exon Length ( bp ) | Intron Length ( bp ) |
| *de novo* | Augustus | 47,723 | 20,378 | 1,148 | 7.6 | 150.9 | 2,911 |
| Homolog | *Danio rerio* | 64,268 | 36,029 | 1,523 | 11.4 | 134.1 | 3,333 |
|  | *Oryzias latipes* | 29,532 | 29,701 | 1,637 | 12.5 | 131.2 | 2,445 |
|  | *Sinocyclocheilus anshuiensis* | 55,080 | 19,982 | 1,644 | 12.6 | 130.3 | 1,579 |
|  | *Sinocyclocheilus rhinoceros* | 57,118 | 21,844 | 1,631 | 12.5 | 130.6 | 1,760 |
|  | *Sinocyclocheilus grahami* | 49,556 | 29,535 | 1,507 | 11.1 | 135.5 | 2,768 |
| Transcriptome |  | 31,114 | 9,515 | 1,628 | 7.6 | 214.7 | 1,198 |
| Consensus | MAKER | 42,205 | 18,601 | 1,331 | 9.0 | 148.2 | 1,996 |

**Note:.** In homolog annotation, we used the genome and gene set of *S. grahami* that were assembled using Illumina data [45].

### Allotetraploid origin of diverse *Sinocyclocheilus* members

To confirm that *Sinocyclocheilus* fishes originated from allotetraploid, we performed a genome-wide synteny analysis of *S. grahami* and *C. carpio* (**Figure 1D-F** and **Figure 2**). Compared with common carp, 18 large chromosomal rearrangements were observed in *S. grahami* genome, including two chromosomal fusions (**Figure 1D**), five chromosomal translocations (**Figure 1E**) and eighteen chromosomal inversions (**Figure 1F**). Among them, chromosome 1 (Chr1) of *S. grahami* was homologous to Chr A22 and A14 of common carp; Chr3 of *S. grahami* was homologous to Chr B22 and A25 of common carp. These alignments made *S. grahami* with two fewer chromosomes than the common carp.

**Figure 2.**
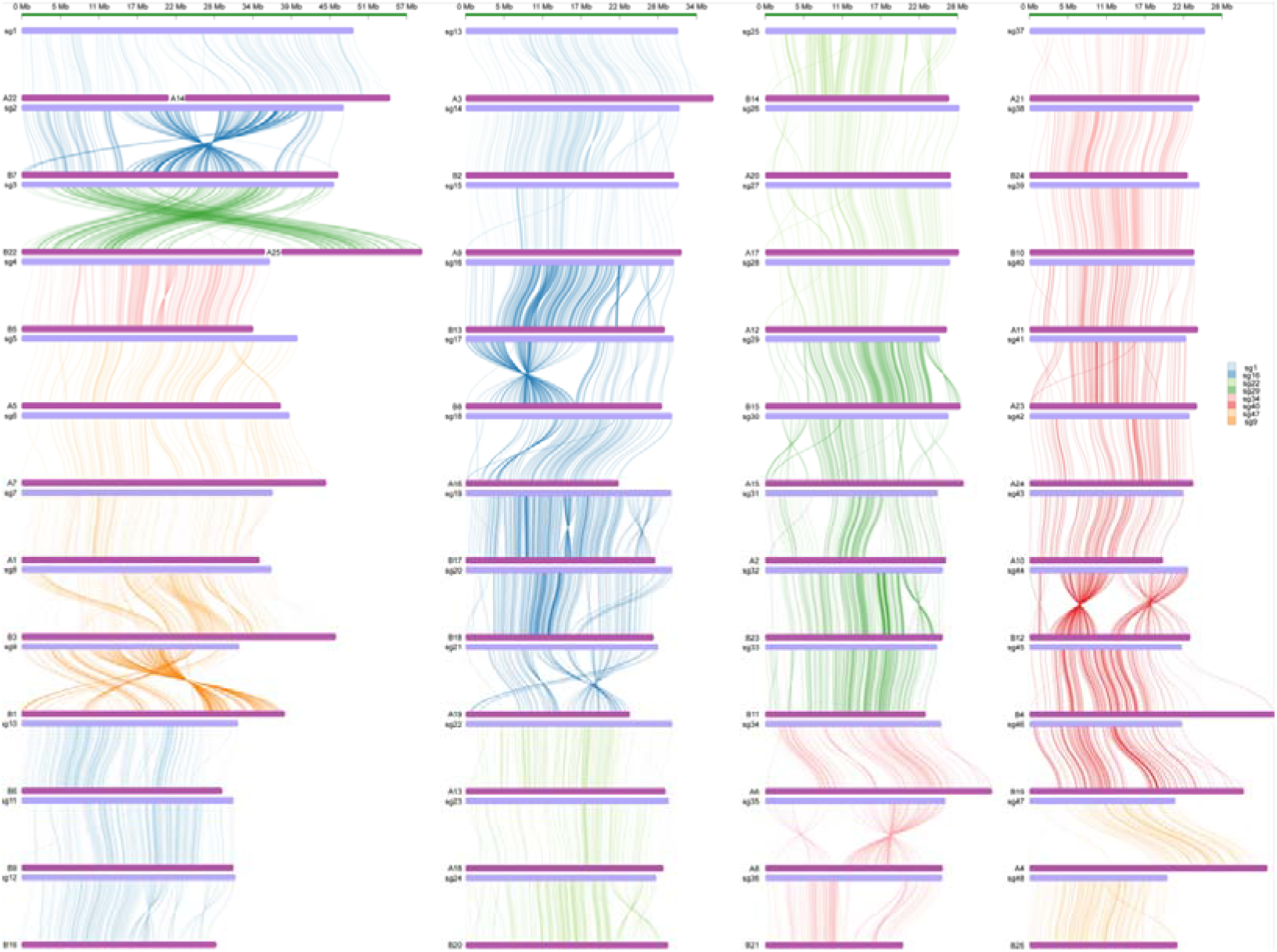
Genome synteny of *S. grahami* (top) and *C. carpio* (bottom)

According to our synteny results, we renumbered the chromosomes of *S. grahami* and divided them into two sub-genomes. The other four *Sinocyclocheilus* members and goldfish were also identified with two sub-genomes for phylogenetic analysis. In the established phylogenetic tree, the group of sub-genome A was clustered into a single branch; the branch of subgenome B was homologous to the ancestors of *O. macrolepis, P. huangchuchieni* and *P. tetrazona* (**Figure 1G**), which is similar to the pattern from early reports [37, 46].

## CONCLUSION

In summary, we constructed chromosome-level genome assemblies for five *Sinocyclocheilus* species. These reference genomics data are valuable genetic resources for in-depth study on phylogenetic evolution and biodiversity of various *Sinocyclocheilus* species, and lay a solid foundation for understanding cave adaption and cavefish biology. Our current study can also contribute to species conservation and exploitation of potential economic and ecological values for diverse *Sinocyclocheilus* members.

## ABBREVIATIONS

Hi-C: high-throughput chromosome conformation capture
LINE: long interspersed nuclear element
LTR: long terminal repeat; Mya, million years ago
SINE: short interspersed nuclear element.

## DATA AVAILABILITY

The genome assembly of *S. grahami* was uploaded to NCBI under the BioPoject PRJNA1172646. All other data, including the repeat and gene annotation, was uploaded to the GigaDB repository, with separate entries for the individual species genomes.

## DECLARATIONS

### Ethics approval and consent to participate

The authors declare that ethical approval was not required for this type of research.

### Authors’ contributions

Q.S. and J.Y. conceived the study and designed the project. Y.O. and J.Y. managed the project and prepared samples. C.B., R.L., Y.O., and X.M. performed data analysis and wrote the manuscript. Q.S. and J.Y. revised the manuscript. All authors contributed to data interpretation.

